# Molecular Scale Spatio-Chemical Control of the Activating-Inhibitory Signal Integration in NK Cells

**DOI:** 10.1101/2020.03.24.004895

**Authors:** Esti Toledo, Guillaume Le Saux, Long Li, Maor Rosenberg, Yossi Keidar, Viraj Bhingardive, Avishay Edri, Uzi Hadad, Carmelo Di Primo, Thierry Buffeteau, Ana-Sunčana Smith, Angel Porgador, Mark Schvartzman

## Abstract

The role of the spatial juxtaposition between activating and inhibitory receptors in cytotoxic lymphocytes has been strongly debated in the context of the inhibition of immune signaling. The challenge in addressing this problem was so far a lack of experimental tools which can simultaneously manipulate different signaling molecules. Here, we circumvent this challenge by introducing a nanoengineered multifunctional cell niche, in which activating and inhibitory ligands are positioned with molecular-scale variability and control, and applied it to elucidate the role of the spatial juxtaposition between ligands for NKG2D and KIR2DL1 – activating and inhibitory receptors in Natural Killer (NK) cells – in KIR2DL1-mediated inhibition of NKG2D signaling. We realized the niche by a nanopatterning of nanodots of different metals with molecular scale registry in one lithographic step, followed by a novel ternary functionalization of the fabricated bi-metallic pattern and its background to with three distinct biochemical moieties. We found, that within the probed range, the 40 nm gap between the activating and inhibitory ligands provided an optimal inhibition condition. Supported by theoretical modeling and simulations we interpret these findings as a consequence of the size and conformational flexibility of the ligands in their spatial interaction. Our findings provide an important insight onto the spatial mechanism of the inhibitory immune checkpoints, whose understanding is both fundamentally important, and essential for the rational design of future immunotherapies. Furthermore, our approach is highly versatile and paves the way to numerous complex molecular platforms aimed at revealing molecular mechanisms through which receptors integrate their signals.

---

Cells communicate with their environment through a rich repertoire of receptors, whose signaling integration is mediated by the biochemistry of the cell-environment interface, as well as by diverse physical features, such as receptor size and spatial arrangement. In the immune synapse – the functional interface between lymphocytes and antigen-presenting cells – activating, costimulatory, and inhibitory receptors coordinate their nanoscale clustering and arrangement to regulate lymphocyte immune activity(1, 2). For example, the spatial density and organization of T Cell Receptors (TCR) and their co-receptors CD3 within their nanoclusters regulate the phosphorylation of their complexes(3). Also, TCR and its linker for activation (LAT) concatenate during T cell activation, but form segregated clusters in resting T cells(4, 5). The inhibitory function of immune checkpoints that orchestrate the self-tolerance of the immune system, such as immunotherapeutic target PD-1 in T cells, or KIR2DL1 in NK cells, is associated with their nanoscale clustering and their segregation from activating and co-stimulatory receptors(6, 7), yet mechanism of these clustering and segregation, and their role in the activating-inhibitory signaling coordination, are still obscure. The systematic study of how the segregation between different receptors regulates their signaling integration requires a high level of their spatial control with respect to each other. However, protocols currently available for cell biologists allow to control receptor segregation only in the direction perpendicular to the membrane, by manipulating the length of their extracellular domains. In the context of immune receptors, controlled variations in the ectodomain size of receptors were shown to affect their signaling integration in T cells and NK cells(8–11). At the same time, there has been no methodology to control segregation of different receptors within the membrane plane, which is more relevant for the regulation of signaling cross-talk in the immune system(12). For this reason, the effect of within-membrane segregation different receptors onto the cell function could not be systematically studied.

Receptors can be spatially guided within the cell membrane by artificial cell niches based on tunable patterns of extracellular ligands. Various concepts for such niches have evolved in the last two decades thanks to the progressing advances in nanofabrication, and they were applied to study the effect of ligand distribution on different cell functions, such as adhesion(13–15), chemokinesis (16), mechanosensing(17), differentiation(18), and immune activity(19–22). Functionalized surfaces were also the basis of mimetic cell models, which were used, in conjunction with theoretical modelling, to identify the physical foundation for the recognition of membrane-confined ligands and receptors, which, mostly focused on single protein pairs(23). However, all of the state-of-the-art ligand patterns, even those including multiple ligands(19, 24), have been limited to spatially control ligands of one type only. Simultaneous control over two or more ligands, on the other hand, has not been realized so far, as it requires new, paradigm shifting fabrication and functionalization approaches with precision and complexity far beyond those exploited for the state-of-the-art ligand patterns.

In this paper, we achieved, for the first time to the best of our knowledge, the molecular-scale spatial control of two extracellular ligands, in order to discover how the spatial juxtaposition between two receptors: (i) NKG2D – the major activating receptor in NK cells, and (ii) KIR2DL1 – an inhibitory receptor that belongs to the family of killer cell immunoglobulin-like receptors (KIRs)(25, 26), regulate KIR2DL1-mediated inhibition of NK cell cytotoxic activity. Remarkably, activating and inhibitory receptors in NK cells balance their signals to determine whether a target cell will be tolerated or attacked. In the context of NKG2D and KIR2DL1, there has been emerging evidence that KIR2DL1-mediated inhibition of NKG2D signaling is closely associated with the nanoscale clustering and arrangement of the two receptors(27), yet the exact role of this arrangement in signaling cross-talk between the two receptors is still unclear. To spatially control both receptors, we realized a molecular-scale device patterned with periodic heterogeneous arrays of their ligands, by selectively anchoring these ligands to nanofabricated arrays of dots of two different metals. We used these arrays as artificial niches for the stimulation of NK cells. While being sized at the scale of individual ligand-receptor complex, the anchoring dots functionalized with ligands guide the clustering of the two receptors, producing, for example, their colocalization (Fig. 1a), or controlled segregation (Fig. 1b), depending on the array geometry. We fabricated the arrays by nanoimprint lithography and double angle evaporation, which allowed to register the anchoring dots of two different metals with molecular-scale accuracy in one lithographic step. To selectively attach the activating and inhibitory ligands to the anchoring nanodots, we developed a novel *ternary* chemical functionalization based on the combination (i) histidine-nitrilotriacetic acid (NTA) conjugation, (ii) biotin-avidin conjugation and (iii) Silane chemisorption (Fig. 1 c), thereby forming, for the first time to the best of our knowledge, three distinct chemical moieties onto a heterogeneous nanopattern.

**Fig. 1.**
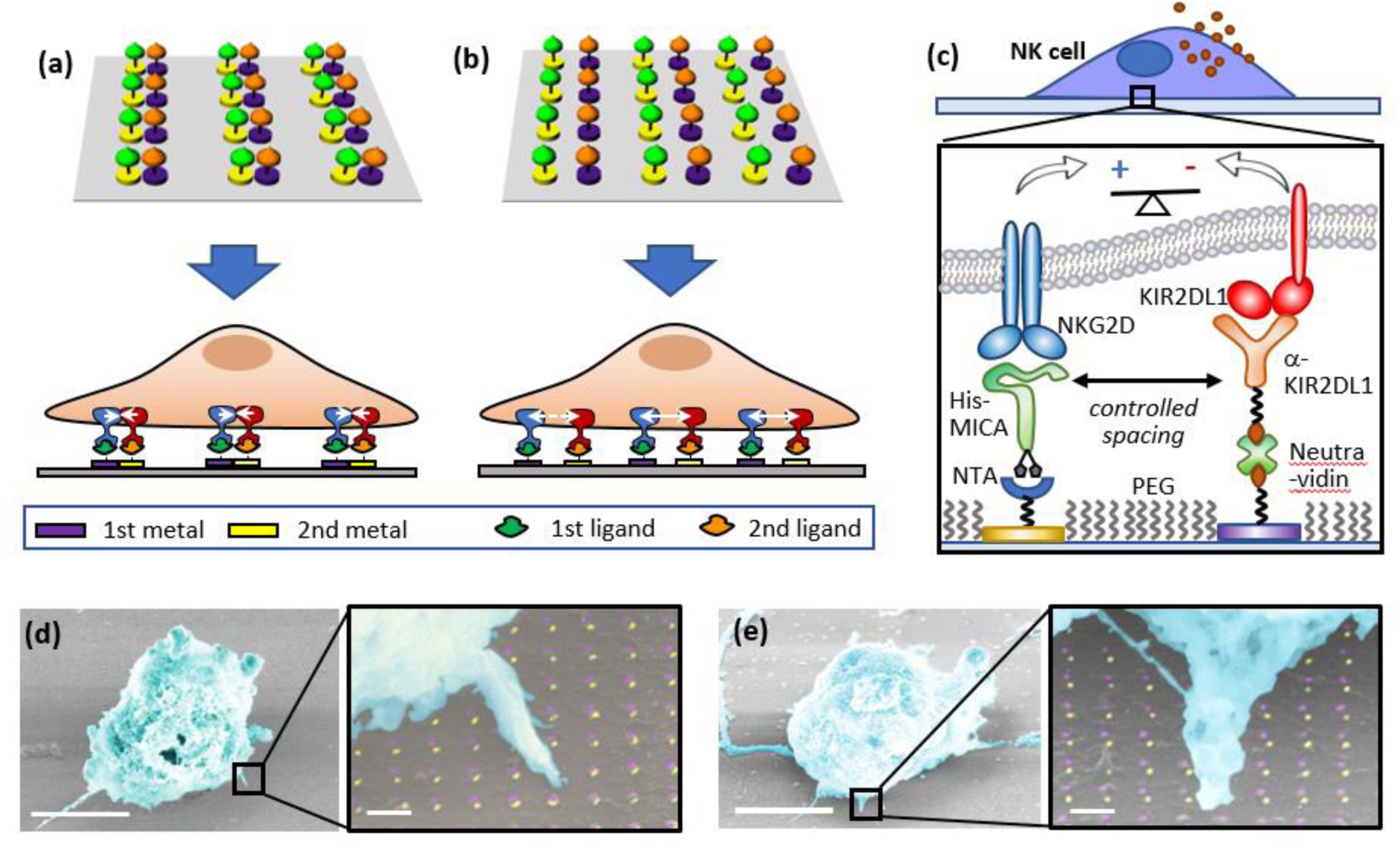
(a) and (b). Schematic representation of ligand patterns in which colocalized or spatially segregated ligands, respectively, control the spatial arrangement of integrating receptors. (c) Scheme of a ligand pattern that control activating-inhibitory balance in NK cells. (d) and (e) Scanning Electron Micrograph of NK cells stimulated in activating-inhibitory ligand array with different spacings between the ligands.

We stimulated NK cells on different arrays with in which only one geometric parameter – the spacing between the two ligands – was systematically varied (Fig. 1 d, e). We found, that NK cells sense this spacing, and adjust their immune response to its extremely small, molecular-scale changes: while colocalized ligands produced an insufficient inhibitory effect on the activation of NK cells, this inhibition gradually increased by inducing a nanoscale gap between the ligands, and became significant at a gap of 40 nm – the largest one within the tested range. This finding indicated the possible role of the conformational flexibility of the ligands in their spatial interaction, and was further rationalized by theoretical modelling, which showed that this spacing corresponds to the optimal separation between the ligands, which stems from the limited flexibility of the receptors and the membrane, and which compensates for the size difference between the studied ligands. This strong correlation between the experimental results and the theoretical model clearly establishes a link between the spatial arrangement of activating and inhibitory receptors and their signaling integration in NK cells, Interestingly, our results show that although signaling of NK cells has been closely associated with nanoscale clustering of activating and inhibitory ligand-receptor complexes in NK immune synapse(27–29), this clustering is not an necessary condition for the inhibition in NK cells, and that the inhibition is also possible for a fixed configuration of controllably distributed ligands, in which the spatial parameters, e.g. ligand spacing, can be used as a tuning knob for the inhibition efficacy. Importantly, besides the specific case study of KIR2DL1-NKG2D signaling cross-talk, our innovative nanodevice technology opens a general pathway to complex, multifunctional nanomaterials designed to model the cell environment with unprecedented precision and complexity, and enable numerous molecular-scale studies of the structure and mechanism of functional biointerfaces, which have been impossible up to date, but are now within reach.

## Device design and fabrication

The device surface was patterned with heterodimers of Au and Ti and nanodots functionalized activating and inhibitory ligands. The heterodimers were configured into orthogonal arrays with the periodicity of 200 nm in both *x* and *y* axis. This periodicity, which is twice larger than the average membrane correlation length(30), came to minimize any possible physical interaction between the neighboring dimers, and at the same time provide sufficient amount of ligands to stimulate NK cells. The gap between the nanodots was the only degree of freedom in the device design, and it varied between 0 to 40 nm. Remarkably, such a delicate registration between two nanofabricated metallic layers is extremely challenging to achieve with standard nanolithographic approaches, especially in a scalable manner. Here, we developed an “out-of-the-box” fabrication approach, in which we combined nanoimprint lithography with double angle evaporation. This combination allowed to produce bimetallic nano arrays registered with nanoscale accuracy using only one lithographic step, and with no need for alignment between different nanodots. Specifically, we first patterned a thermal nanoimprint resist on a Si substrate with periodic rectangular arrays of ∼ 20 nm holes separated by 200 nm, and transferred the imprinted pattern into a resist by angle deposition of a metallic protection mask(31), resist over-etch by plasma, two sequential metal evaporations (Au and Ti/Cr in our case), and lift-off (Fig. 2a). The resulting nanopattern consisted of bi-metallic heterodimers, whose rectangular arrangement and 200 nm periodicity were determined by the initial nanoimprint pattern. The size of each nanodot was 15-20 nm, and the gap between the two neighboring nanodots within the dimers was determined by the evaporation angles (Fig. S1). In such a way, small evaporation angles produce heterodimers of colocalized nanodots (Fig. 2b), whereas a gradual increase in the evaporation angle resulted in a gradual separation between nanodots (Fig. 2b - e).

**Fig. 2.**
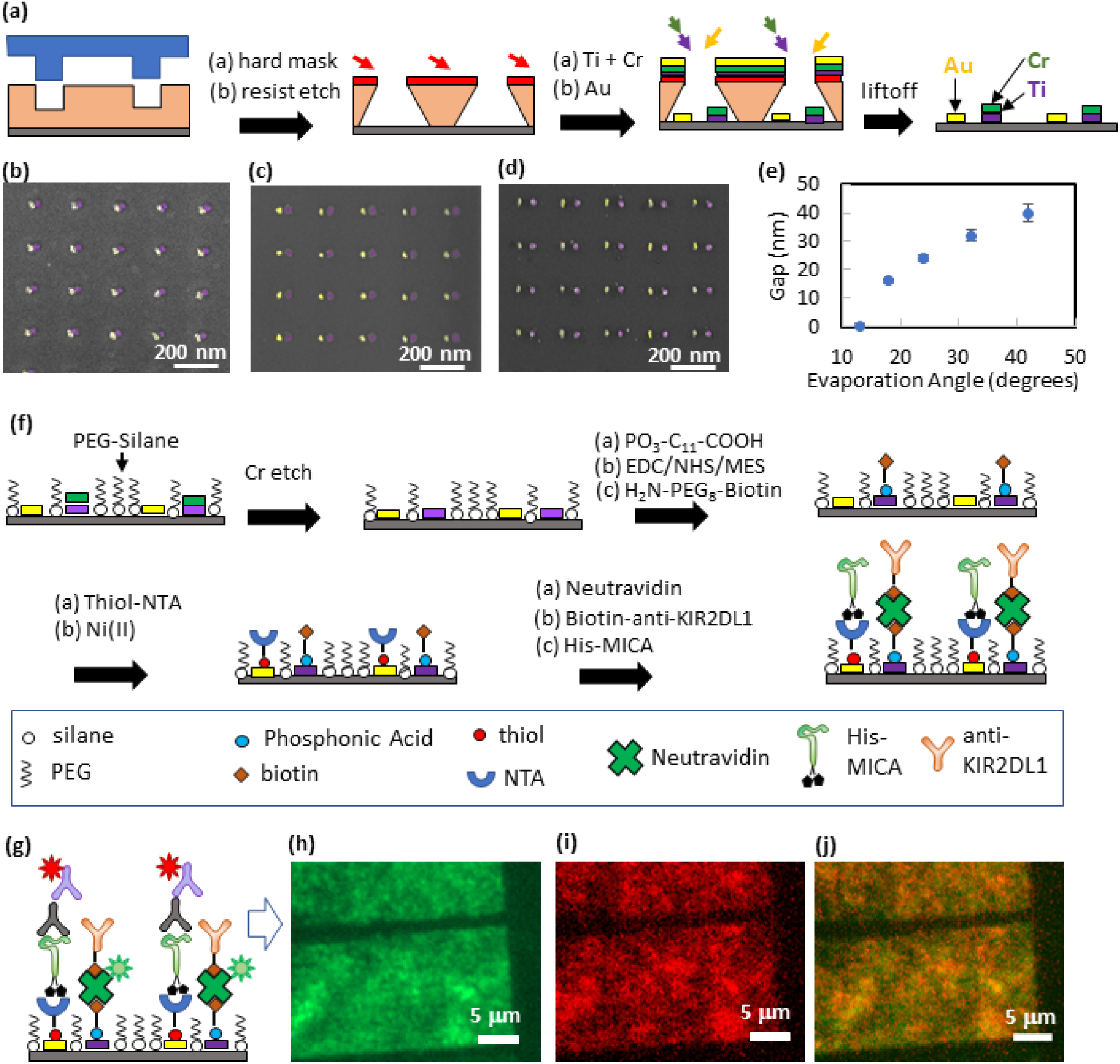
(a) Fabrication of heterogeneous metallic arrays. (b) Colocalized nanodots (c) Nanodots separated by ∼ 20 nm gap (d) Nanodots separated by ∼ 40 nm gap. (e) Segregation control via evaporation angle (f) Functionalization scheme (g) Scheme of the fluorescent labelling of the functionalized nanodots. (h)-(j) Fluorescent imaging of a functionalized array taken in green channel, red channel, and their merge, respectively.

The second step in the device fabrication is the biofunctionalization of the bi-metallic nanoarrays with the desired ligands. The key challenge here is to achieve the absolute site-selectivity with which different ligands are immobilized onto different nanodots. Remarkably, not only each of the two used ligands should be specifically immobilized on the selected nanodots, but also neither of these two ligands is allowed to attach to the Si background surface that surround the anchoring nanodots. To achieve this, as well as to prevent any non-specific interaction between the background and the cell receptors, the Si background should be functionalized with an antifouling agent. Therefore, the surface of our nanodevice includes *three* distinct and discretized chemical entities – the background and two ligated nanodots. To date, state-of-the-art biomimetic patterns for controlled cellular environment consisted of up to two spatially segregated biochemical functionalities, i.e. one ligand and antifouling agent(32), or two ligands(19). Here, we developed a novel *ternary* functionalization process for the site-specific immobilization of three biochemical functionalities – two ligands and an antifouling agent, by combining chemical modifications of Si, Au, and TiO_2_ surfaces with organic silanes, thiols, and phosphonic acids, respectively (Fig. 2f). First, the Si background was passivated with silane terminated polyethylene glycol (PEG) – a commonly used antifouling agent that effectively prevents non-specific binding interactions(33, 34). Importantly, the unwanted immobilization of the silane onto oxidized Ti nanodots was prevented by covering them with a protective Cr layer during the fabrication process. This Cr layer was etched after Si passivation with PEG. The exposed oxidized Ti was then functionalized which a carboxyl-terminated phosphonic acid, and subsequently modified with a biotin terminal group, while PEG-Silane effectively protected Si from the non-specific chemisorption of phosphonic acid. Then, Au was functionalized with an organic thiol terminated with nitrilotriacetic acid (NTA), which was next chelated with Ni ions to provide high affinity to the Histidine conjugate. (See supporting information and Figs. S5-7 for PM-IRRAS analyses that confirm the chemical modification of Au and TiO2). Finally, His-tagged MICA was attached onto Au nanodots via His-NTA/Ni conjugation, and biotin labelled anti-KIR2DL1 was attached onto biotin functionalized TiO_2_ nanodots *via* Neutravidin bridge. Importantly, we used here a monoclonal antibody for KIR2DL1 that was shown not only bind KIR2DL1 receptor, but also induce KIR2DL1-mediated inhibition(35), thus we created here a surrogate antigen presenting cells with fully functional activating and inhibitory stimulating features for NK cell.

To verify the functionalization efficiency and selectivity, we imaged the functionalized array by fluorescence microscopy. Specifically, we verified TiO_2_ functionalization by imaging Neutravidin labelled with Oregon Green 488. In parallel, we verified Au functionalization by immunostaining immobilized MICA with mouse anti-MICA antibody and anti-mouse Alexafluor 568 (Fig. 2g). Fig. 2 h and i show fluorescent images of the functionalized array taken separately in the green and red channels, which represent each of the immobilized bio-functionalities. Also, Fig. 2 j shows the merged image of the two channels. Remarkably, the borders of the array areas are clearly distinguishable in all three images, as the surrounding background produces little fluorescence, and therefore confirms that the ligands were immobilized specifically onto the nanodots. We must note that our ternary functionalization approach was demonstrated here on a specific set of ligands. However, since NTA/Ni-His and biotin-avidin links are very commonly used bioconjugation techniques, and the variety of biomolecules that can be tagged with either biotin or histidine is practically unlimited, our ternary approach is very versatile, and can be used as a general route to produce multifunctional molecular-scale ligand patterns.

Finally, we verified that our device could control ligand arrangement at the single molecule scale. It is generally hypothesized that when biomolecules are immobilized onto lithographically patterned nano-features, their occupancy is determined by steric hindrance(13, 14). Recently, this hypothesis was recently confirmed experimentally using fluorescence microscopy(22, 36). In our devices, we functionalized the nanodots with MICA, whose complex with NKG2D has a footprint of 12 x12 nm (37), and anti-KIR2DL1, whose footprint is about 7 × 19 nm (38). Hence, the size of the fabricated nanodots suits the size of the immobilized molecules, so that each nanodot hosts on average one to two ligand-receptor complexes. To further confirm this, we immobilized KIR2DL1 and MICA onto metallic films according to our functionalization protocol, and performed surface plasmon resonance (SPR) analyses, which showed that the footprint of both biomolecules is about 200 nm^2^ each (see SI and Figs. S2-4 for details).

## Activation of NK Cells

With the demonstrated nanoscale control over the spatial segregation of different ligands, we could now explore how the segregation between ligands for NKG2D and KIR2DL1 receptors affects KIR2DL1-mediated inhibition of NKG2D signaling in NK cells. For this purpose, we stimulated primary NK cells on arrays of MICA and anti-KIR2DL1, which were either colocalized, or separated by gaps of 20 nm and 40 nm. We also used three types of control samples: (i) positive control samples that contained MICA immobilized on a continuous Au film, and should provide optimal conditions for the dense clustering of NKG2D, and thus maximize the activation of NK cells, (ii) negative control samples that contained a continuous TiO_2_ film functionalized with anti-KIR2DL1, which was assumed to provide the optimal condition for the dense clustering of KIR2DL1 within the cell membrane, and (iii) neutral control samples of glass surface covered with poly-l-lysine, which enhances cell adhesion but lacks any specific functionalities for immune stimulation.

We assessed the cytotoxic activity of NK cells by monitoring the fluorescent intensity of lysosome-associated membrane protein-1 CD107a (39, 40) (Fig. 3a), which is a broadly used marker for the immune activation of T cells and NK cells(41, 42). In activated NK cells, lytic granules that contain CD107a move to the immune synapse, fuse with the cell membrane, and degranulate, thereby exposing CD107a. In our case, quantifying the expression of CD107a is the ultimate way to assess the degree of the immune activation of NK cells, since the resulting biological information obtained by other methods like western blot or flow cytometry would stem from cells activated on and outside of the ligand pattern. Fig. 3 b and c presents representative images of NK cells stimulated on patterned and control surfaces, showing the expression of CD107a (white colored).

**Fig. 3.**
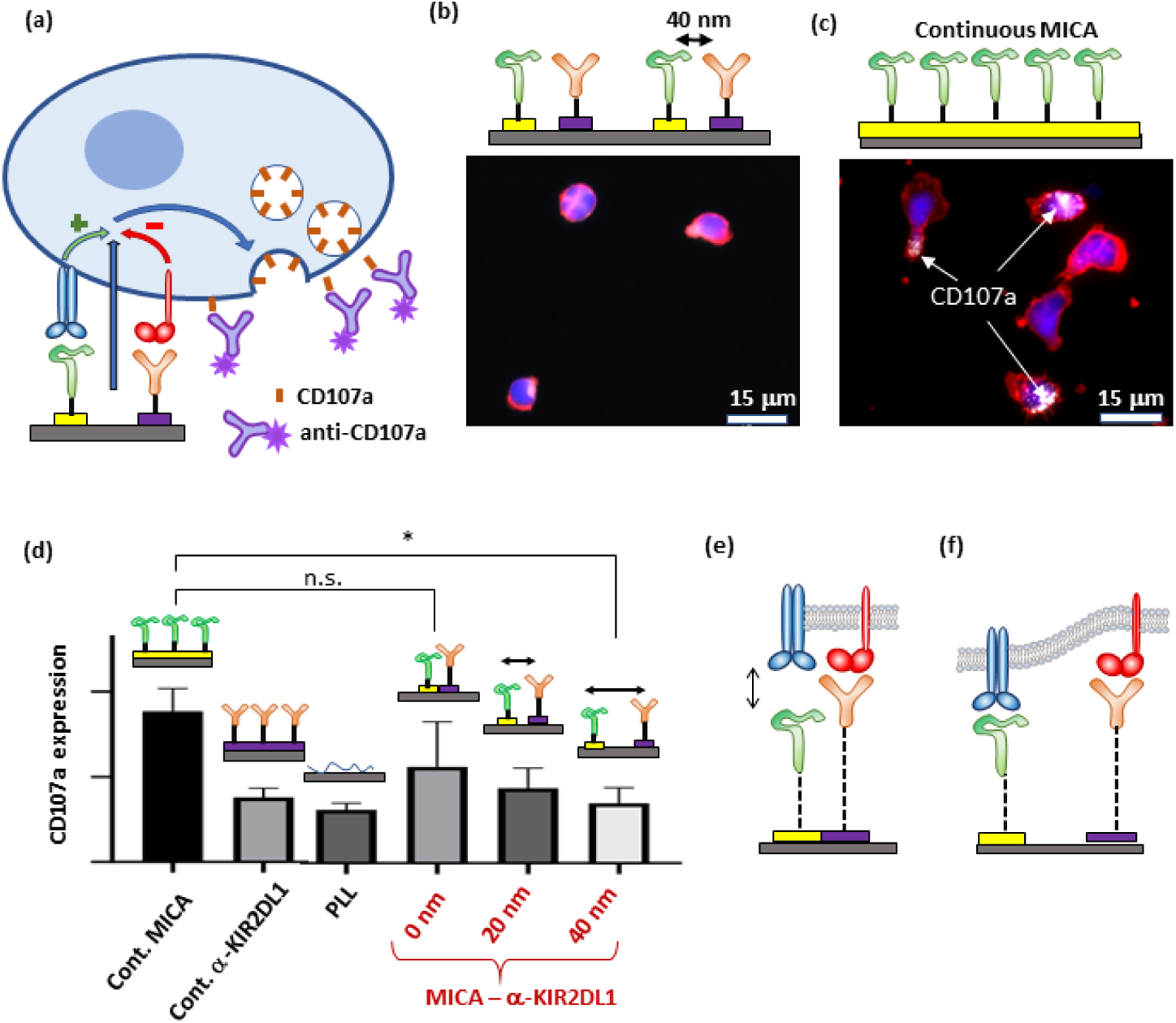
(a) Scheme of CD107a expression regulated by the spatial balance between activating and inhibitory ligands. (b) and (c) NK cells stimulated on nanopattern of MICA and anti-NKG2D (gap of 40 nm), and continuous MICA film, respectively. The cells were stained with phalloidin for cytoskeleton (red), DAPI for nucleus (blue), and anti-CD107a (white). (d) Average amount of CD107a per cells on ligand patterns compared to that in cells stimulated on control surfaces. Surfaces with continuous MICA provided the optimal condition for the activation of NK cells, whereas surfaces with PLL and KIR2DL1 did not stimulate significant CD107a expression due to the absence of activating ligands. Patterned arrays of MICA and anti-KIR2DL1 produced activation that depended on the gap between the ligands. (e - f) Schemes of the possible mechanisms explaining the observed effect of the gap between the ligands in NK cell activation: (e) When both the ligands are colocalized, NKG2D is not engaged by MICA due to the difference in the ligand spacer height, (f) Both receptors are engaged when separated from each other due to membrane flexibility.

As expected, the continuous MICA film provided the optimal conditions for the immune activation NK cells (Fig. 3d). This observation mirrors recent studies that employed diverse microscale and nanoscale stimulating platforms to reveal the effect of clustering and density of activating ligands on the immune response of T cells and NK cells, (20, 21, 43). It is reasonable to assume that the continuous MICA film used in our experiments provides a much higher amount of MICA ligands than the threshold amount required for the activation of NK cells, thus the obtained average CD107a signal was relatively high. On the other hand, NK cells stimulated on poly-l-lysine and continuous anti-KIR2DL1 expressed on average similar amounts of CD107a, which are almost one third of that found for the MICA surface. Specifically, low levels of CD107a expression by the cells stimulated on anti-KIR2DL1 surface can be explained by the lack of any activating functionality rather than by the presence of anti-KIR2DL1 itself. Indeed, inhibitory signaling of KIR2DL1 suppresses NKG2D-mediated activation by blocking NKG2D clustering within the immune synapse (44), thus the recognition of KIR2DL1 itself is unlikely to produce any observable effect on the immune response of NK cells.

The most intriguing result, however, is the effect of the gap between anti-KIR2DL1 and MICA on the activation of NK cells. (Fig. 4d). NK cells stimulated on arrays with colocalized ligands produced, on average, a CD107a signal that was slightly lower than that obtained continuous MICA, yet not significantly. This result indicates that when anti-KIR2DL1 colocalizes with MICA, it cannot not produce a sufficient inhibition of the NKG2D signal. The inhibitory effect, however, progressively increased with the gap between the two ligands, and became significant for the gap of 40 nm. This result demonstrates that the nanoscale architecture of activating and inhibitory ligands can encode the immune response of NK cells. Indeed, regardless of ligand spacing, ligand density is constant for all the tested activating/inhibitory conditions. Furthermore, the projected cell areas are very similar on all surfaces, suggesting that cells are exposed to the same total number of ligands of both types. Therefore, the observed pronounced effect on the inhibitory-activating balance in NK cells was obtained solely by altering the nanometric gap between the two ligands, while keeping all other compositional and spatial parameters of the cell environment constant.

**Fig. 4.**
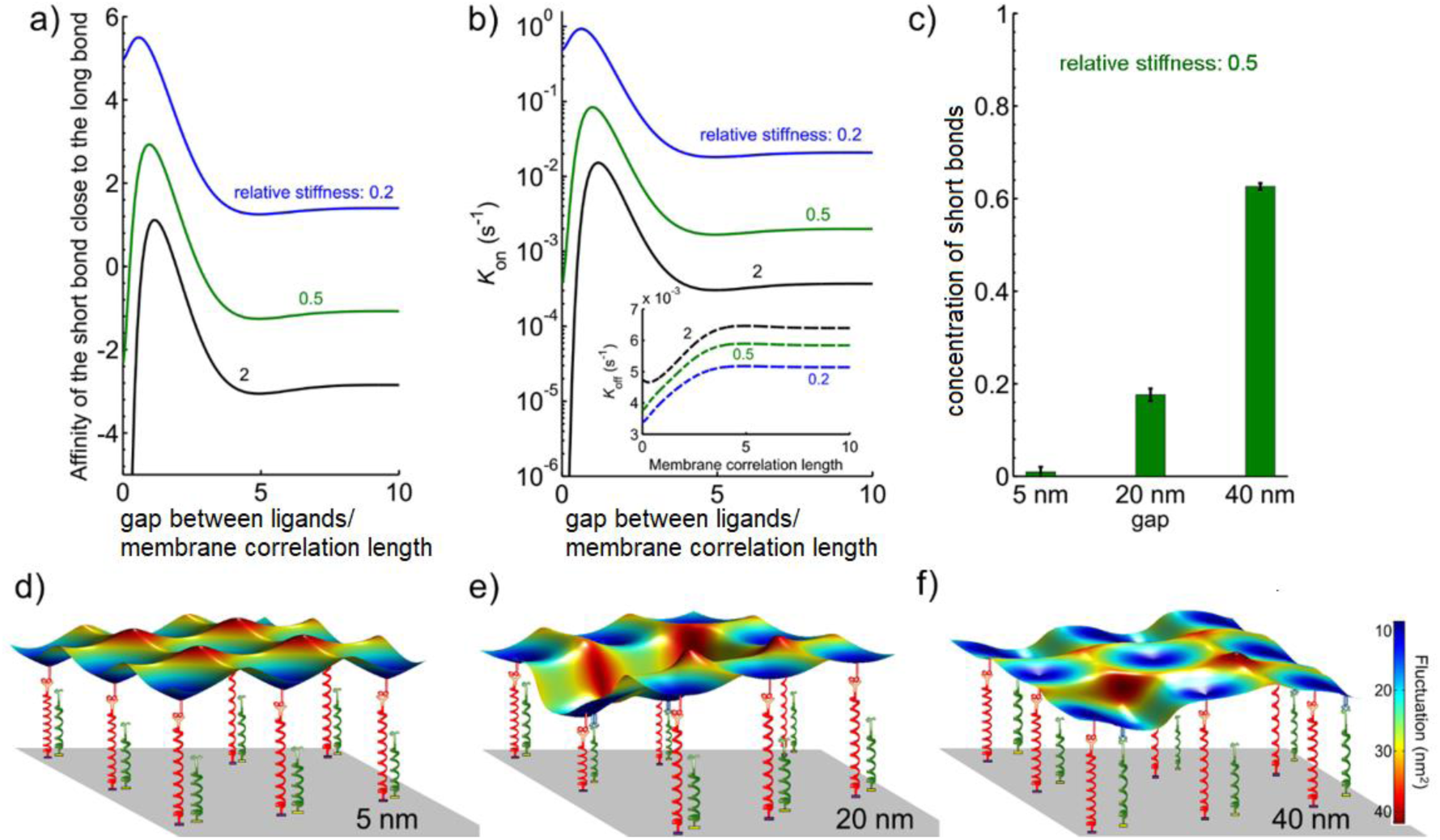
Modelling the interaction of a NK cell with a substrate with immobilized ligands for KIR2DL1 and NKG2D. a) Affinity and b) binding rates of the short bond as a function of a distance (normalized to the membrane correlation length which is about 100 nm) from the existent long bond. c) Concentration of total number of bonds for different gap sizes, showing a clear increase in their numbers when the gap is wider. Characteristic snapshots of a small segment between the membrane and the patterned substrate is shown for a gap of d) 5 nm, e) 20 nm and f) 40 nm.

The observed effect of the ligand segregation onto KIR2DL1-mediated inhibition is not obvious. We hypothesize that this effect is related to the difference in the total length of the ligand constructs (including the ligand molecule and the linker to the nanodots) and the relative flexibility of the bonds. In our case, the total length of MICA, including thiol-NTA linking molecules, is about 10 nm, while the total length of the anti-KIR2DL1 mAb including its biotin-avidin attachment chemistry on Ti is about 22 nm (See SI for the estimation of the ligand length). For colocalized ligands, this length difference results in a ∼13 nm mismatch between the ligands in the direction perpendicular to the membrane. Such a mismatch could prevent the simultaneous engagement of adjacent KIR2DL1 and NKG2D receptors, as schematically shown in Fig. 3e, which could then explain how KIR2DL1 mediated inhibition is impeded (11). On the other hand, both receptors can be engaged when the ligands are spaced by a small gap within the membrane plane, depending on the flexibility of the membrane and receptor-ligand complexes (Fig 3f).

## Modeling of ligand-receptor the interactions

To verify our hypothesis, we adapted the recently developed model that quantitatively captures interaction rates of ligand-receptor pairs embedded in membranes(45) (see SI, section 9, for details). This model is particularly suitable here, since no direct involvement of the cytoskeleton is expected in the formation or dissociation of KIR2DL1-anti-KIR2DL1 and NKG2D-MICA complexes. Accordingly, we mathematically represented each bond as a fluctuating flexible spring with a defined experimental rest length and stiffness. Consequently, we calculated the affinity of the shorter bond (Fig. 4a) from its binding and dissociation rates (Fig. 4b) in the presence of the longer bond, and vice versa (Fig. S9). Notably, the rates themselves are obtained from the convolution of the membrane gap-dependent distribution of positional probabilities with Bell-Dembo-like kinetics(46) (the dissociation rate is dependent on the force acting on the bond, while the binding rate reflects the energy stored in the bond). In the case of an isolated pair, consisting of one short (activating) and one long (inhibitory) bond in the membrane, these rates show a non-monotonous behavior, and both bonds in the pair are stabilized relative to the lone bond. Furthermore, there is an optimum distance of about half of the membrane correlation length, at which each receptor has the highest binding rate and affinity for its cognate ligand. This phenomenon stems from the fact that the bonds are of different lengths and is further promoted by the difference in bond flexibilities. Importantly, membrane correlation length is of the order of 100 nm, thus the modelled optimal spacing precisely fits the experimentally observed gap of 40 nm between the ligands(30), at which the inhibitory effect on the stimulation of NK cells was significant.

To verify if this sensitivity to both the mechanical properties of the membrane and to length mismatch persists in a geometry of 2D arrays of immobilized ligands, we performed effective Monte Carlo simulations(45) (see SI, section 10, for details). Specifically, the experimental patterns are first reproduced in our *in silico* model of the substrate, with respect to both the lateral distribution and the in size of the ligand constructs. Next, the NK cell mimic is allowed to hover over the patterned substrate and spontaneously make and break KIR2DL1-anti-KIR2DL1 and NKG2D-MICA in the contact area with the substrate, using the same rates as discussed above. The number of formed bods in recorded as a function of time and system is allowed to propagate until the steady state is achieved, and the average number of formed bonds is reported (Fig. 4c) Strikingly, we find that similarly to the 2-bond case, at equilibrium, the total number of bonds increases when the gap between the nanodots on the pattern is widened within the range observed experimentally (Fig. 4c). More specifically, the number of short bonds increases with ligand spacing while the number of long bonds is barely affected (Fig. 4d-f).

## Discussions and conclusions

Lymphocyte activation is controlled by numerous receptors, whose spatial arrangement with respect to each other has been hypothesized to depend on their matching sizes : similarly-sized ligand-receptor complexes come in close proximity, whereas differently sized ones remain spatially separated (47). This size dependent separation is the basis of the kinetic segregation model for T cell activation, by which large molecules of tyrosine phosphatase CD45 are spatially excluded from relatively short TCR-MHC complexes formed at the T cell – APC interface, thereby diminishing CD45 mediated dephosphorylation of TCR-MHC, and producing downstream signaling that finally leads to the activation of T cells (48). The hypothesis that the size mismatch between TCR and CD45 causes their segregation upon TCR engagement, and the subsequent decrease in TCR-MHC dephosphorylation, was directly confirmed using several artificial model systems for T cell stimulation. For example, nanofabricated mono-ligand patterns were used to demonstrate that MHC ligands positioned on nanoparticles within the supported bilayer do not allow TCR association with the diffusing CD45, while MHC ligands elevated on etched nano-pillars increased the spacing between the bound membranes and allowed the association of CD45 and TCR, thereby affecting the phosphorylation of the TCR, and modulating T cell immune response (19). Also, a T cell mimicking model based on membrane vesicles was used to show that large CD45 isoforms were excluded more rapidly from TCR-MHC complex than a smaller ones, directly indicating the crucial role of CD45 size in this exclusion (49). Furthermore, elongation of TCR ligands(8) and shortening of CD45 ectodomain (9) and were both shown to suppress TCR engagement.

Whilst T cell immune function is regulated mostly by TCR, the immune function of NK cells is regulated by multiple receptors with different functionalities - activating, costimulatory and inhibitory – whose delicate signaling balance determines the immune response of NK cell. Although receptor clustering in NK cells has been less studied than for T cells, it would be reasonable to presume that the association or segregation of two different receptors in NK cells is regulated by spatial exclusion: two receptors will spatially associate only if the sizes of their complex with their cognate ligands match, and segregate if the sizes are different, similarly to how TCR-MHC and CD45 in T cells do. Indeed, it was previously shown that varying ectodomain size of either MICA or KIR2DL1 ligand HLA-C could greatly affect the inhibition of NK cells: differently sized ectodomains of MICA and HLA-C produced segregation between the two ligands, resulting in a reduction of KIR2DL1 mediated inhibition whether the longer bond was MICA or HLA-C. In contrast, matched ectodomain lengths, regardless of their absolute size, produced colocalization of the two ligands, which allows for both receptors to be engaged and therefore induce an effective inhibition (11).

It should be noted, that in all of the above-mentioned works, the coordinates of two signaling molecules were varied with respect to each other on the axis perpendicular to the membrane, yet, the receptors could still freely move within the membrane plane, with no external control over their colocalization or segregation. Our experimental approach is fundamentally different: we designed and realized a smart system that allows full spatial control of the ligands in all the three dimensions. Varying the coordinates of each ligand in a systematic and independent way allows to methodically study the interplay between ectodomain length and within-membrane segregation of the two ligands, and the effect of this interplay on the signal integration of the two receptors.

In our experiments, we mismatched the coordinates of NKG2D and KIR2DL1 ligands on the axis perpendicular to the membrane, via differently sized ligand spacers. These mismatched coordinates produced conditions for which both ligands, while laterally colocalized within the membrane plane, cannot simultaneously engage their cognate receptors. Thus, we did not observe any inhibition effect for the laterally colocalized ligands. This result largely mirrors the previously reported negative effected of mismatched ligand ectodomains on KIR2DL1 mediated inhibition(11). On the other hand, we experimentally demonstrated that this mismatch of the two ligands on the axis perpendicular to the membrane can be compensated by their appropriate lateral segregation within the membrane plane, resulting in both receptors being engaged. Furthermore, we provided theoretical support for this demonstration based on the modelling of the membrane and receptor conformation. Thus, our work clearly confirms that the high level of proximity of inhibitory and activating ligand-receptor complexes, or tightly packed segregated clusters, are in fact deleterious to inhibitory signaling. Our results furthermore point to the important aspect of biomechanical optimization of the membrane and the protein design in NK cells, in the context of length and stiffness.

We also cannot exclude that the spatial manipulation of two ligands could affect the recruitment of tyrosine phosphatases SHP-1 and SHP-2, which are key players in the inhibition mechanism in NK cells. SHP-1 and SHP-2 are recruited by KIR-family inhibitory receptors, and this recruitment causes tyrosine dephosphorylation of activating receptors such as 2B4, NKG2D, and their costimulatory receptors (50, 51). Whether this mechanism requires the physical recruitment of SHP-1 and SHP-2 between the activating and inhibitory receptors is unclear. If this is the case, co-localizing MICA and anti-KIR2DL1 by their immobilization onto overlapped nanodots (with a zero gap) could prevent the recruitment of SHP-1 and SHP-2 in the interstice between the two receptors, which might be needed for the effective tyrosine phosphorylation of NKG2D. It remains uncertain, however, if a ligand spacing of a few tens of nanometers, as in our most potent inhibitory system, can impede the association between NKG2D and KIR2DL. Given that the successful regulation of NK cell activation has been observed in segregated domains, the role of the direct interactions between NKG2D and KIR2DL, and its impact on KIR2DL1 inhibition remains an open question.

In summary, we demonstrated here that controlling the juxtaposition of activating and inhibitory ligands either on the axis perpendicular to the membrane plane, or within the membrane plane, individually or in combination with each other, provides a powerful means to regulate the functional cross-talk in NK cell activation. We trace the potential origins of this behavior in the biomechanical optimization of the receptor-ligand constructs and the membrane. This exquisite control is only possible by a precise lateral fixation of ligands, which was allowed by our novel lithographic approach combined with unprecedented site-specificity of ligand immobilization intro three distinct functionalities. In addition to the specific case study of the inhibitory-activating signal integration in NK cells, the nanotechnological platform supported by the theoretical insights, allows broad studies of the role of spatial coordination of different signaling molecules in their functional interplay. Such an interplay could be important for the activation of other immune cells, such as T cells and B cells, whose signaling mechanism is yet to be fully understood. Beyond the fundamental insights into the immune cell regulation, our approach could be useful in more applied context of immunotherapy. One example could involve systems based on dual activating and inhibitory Chimeric Antigen Receptor (aCAR:iCAR)(52), whose efficient delivery requires the optimization of a combination of multiple ligated receptors. The work presented here establishes a reliable methodology for comprehensively addressing various systems, in which the integration of multiple signals from multiple receptors needs to be identified and optimized.

## Methods

### (Detailed experimental protocols are provided in Supporting Information)

#### Preparation of the bimetallic nanodot arrays

Nanoimprint mold was prepared by electron beam lithography of Hydrogen Silesquioxane (HSQ, XR-1541), and consisted of orthogonal array of ∼20 nm features separated by 200 nm. Thermal resist was imprinted onto Silicon substrates, covered by Titanium mask using angle evaporation, and etched with Oxygen plasma. Ti/Au (1 nm/ 5 nm) and Ti/Cr (5 nm/ 3 nm) were the sequentially evaporated at two opposite angles to produce arrays of nanodot heterodimers after the liftoff.

#### Ternary biofunctionalization of nanodot arrays

Fabricated arrays were cleaned in organic solvents and Oxygen plasma, and Si background was passivated by immersing the samples 48 hrs. in a toluene solution containing 1:10000 w/v PEG-silane and 0.1% v/v acetic acid. Then, Cr protection layer was etched in commercial Cr etchant. Oxydized Ti was biotinylated by immersing samples into the solution of PO_3_–C_11_–COOH for 24 h, baking at 120 °C for 24 h, and overnight incubation in aqueous solution of NH_2_–PEG_8_–biotin. Au was functionalized by reacting overnight in ethanolic solution of thiol–NTA. The samples were blocked for 30 min at 37 °C in PBS with 5% skim milk, followed by 30 min incubation in 25 μg/mL NeutrAvidin Oregon Green 488 in PBS with 5% skim milk. The samples were then immersed overnight in a solution containing 2 μg/mL of biotin α-KIR2DL1 in PBS with 5% skim milk. NTA chelation with Ni was done by the incubation of the substrates in nickel(II) chloride, followed by rinsing in water and incubation in His-MICA solution in PBS. For cell studies, freshly prepared samples were used as is. To confirm the selective attachment of MICA by fluorescence, the surfaces were first incubated overnight in PBS solution of α-MICA for 1 hr., and then in PBS-solution of anti-mouse Alexa Fluor 555. Finally, surfaces were rinsed once in water, mounted with Dako Fluorescent Mounting Medium and imaged with fluorescent microscope.

#### Preparation of the nanoarrays with varying metal ratios

Silicon substrate with Silicon Oxide layer of 100 nm was patterned by electron beam lithography using positive resist. The pattern consisted of multiple arrays of rectangles (50 nm x 20 nm) oriented at different angles. 20 nm of Silicon Oxide was the etched in diluted Buffered Oxide Etchant (BOE), and Ti/Au (5 nm/ 15nm) and Ti/Cr (15nm/10nm) were sequentially evaporated while the substrates were tilted by 22° and rotated by 90 between the evaporations, following by liftoff. Using a simulation by PVSyst6.0.1 and linear optics calculations using MATLAB, evaporating parameters were calculated so the feature orientation will encode a predefined Au/ Ti ratio in the fabricated rectangles, and SEM of the fabricated arrays were used to confirm the simulated ratios (see Supporting Information).

#### Primary NK Cell Purification

pNK cells were purified from peripheral blood of healthy, adult, volunteer donors, recruited by written informed consent, as approved by the Institutional Review Board Ben-Gurion University of the Negev. The cells were isolated using a human negative selection-based NK isolation kit (RosetteSep, Miltenyi Biotec). The purified NK cells were then cultured in a stem cell serum-free growth medium (CellGenix GMP SCGM, 20802-0500) supplemented with 10% heat-inactivated human AB plasma from healthy donors (Sigma, male AB, H-4522), 1% l-glutamine, 1% Pen-Strep, 1% sodium pyruvate, 1% MEM-Eagle, 1% HEPES 1 M, and 300 IU/mL recombinant human IL-2 (PeproTech).

#### NK Cell Activation Studies

Cultured pNK cells were seeded onto the surfaces in growth medium containing <2% serum, 50 units of Il-2 which was supplemented with APC α-CD10a (1:1000 v/v) and left to adhere for 3–4 h. The surfaces were then rinsed twice in PBS to remove the nonadherent cells, followed by fixing the adherent cells with 4% paraformaldehyde (PFA) and then direct staining with Alexa Fluor 555 phalloidin without permeabilization to prevent damage to the cell membrane. Finally, the nuclei were stained by mounting the samples with ProLong Gold antifade reagent containing DAPI (both from Life Technologies).

We stained CD107a with its fluorescent antibody while the cells were incubated for three hours and then fixed. To prevent the diffusion of CD107a antibody into the cell cytosol and ensure that only CD107a immobilized the cell surface is imaged, the cells were not permeabilized after fixation.

#### Microscopy

The characterization of the surfaces as well as NK cell adhesion and activation were performed using a Nikon Ti2e epifluorescence microscope and quantified using the Fiji imaging software (https://fiji.sc). For α-CD107a quantification of fluorescence intensity, exposure time, and magnification were not changed between samples. We quantified the degree of activation, which is proportional to the intensity of the α-CD107a signal (42), the degree of NK cell activation can also be quantified.

#### Statistics

At least 10 fields at 20× magnification on each surface were analyzed. The data were averaged for each experiment, and the experiments were performed three times. Statistical analysis was performed by analysis of variance, and Tukey’s multiple comparison *post hoc* test was also performed using the Prism software (GraphPad Software Inc., USA). The results were considered to be significantly different for p < 0.05.

## Acknowledgements

- This work was funded by the Multidisciplinary Research Grant – The Faculty of Health Science in Ben-Gurion University of the Negev, Israel Science Foundation, Individual Grant # 1401/15, and Israel Science Foundations: F.I.R.S.T. Individual Grant # 2058/18. SPR and PM-IRRAS analyses were performed with the financial help of COST Action CA15126. This work has benefited from the facilities and expertises of the Biophysical and Structural Chemistry platform at IECB, CNRS UMS3033, Inserm US001, Bordeaux University. The Biacore T200 instrument was acquired with the support of the Conseil Régional d’Aquitaine, the GIS-IBiSA, and the Cellule Hôtels à Projets of the CNRS. We thank Lætitia Minder, technical manager of the SPR Platform. ASS and LL thank the German Science Foundation project SM 289/8-1, AOBJ: 652939. LL is supported by the China Scholarship Council (CSC, File No. 201806185038). ET is supported by Israel Ministry of Science and Technology, Ariane de Rothschild Women’s Doctoral scholarships program for outstanding female Ph.D. students, and Israel Scholarship Education Foundation (ISEF).

