## Supporting Info for "Molecular Scale Spatio-Chemical Control of the Activating-Inhibitory Signal Integration in NK Cells"

### Supporting Information

#### 1) Production of nanodot heterodimers and bimetallic nano-islands

*Preparation of the nanodot heterodimers*

The fabrication of the arrays of nanodot heterodimers is based on nanoimprint lithography (1–3). First, we produced a nanoimprint mold using electron beam lithography of hydrogen silsesquioxane (HSQ) , a negative tone resist. For this, a silicon substrate(<100> orientation, P-type, 5-10 Ω.cm, thickness 380±25 µm) was cleaned with hydrofluoric acid to remove the native oxide layer. The surface was then spin-coated with 20nm of HSQ (XR-1541, Dow Corning) diluted in Methyl Isobutyl Ketone. The HSQ film was patterned by Electron beam lithography using a Raith e-LINE system and consisted of orthogonal arrays of ~20 nm features separated by a distance of 200 nm. The writing parameters were as follows: current 190 pA, aperture 30µm, acceleration voltage 10 kV. The patterned substrate was then developed in AZ 726 MIF Developer (Microchemicals), rinsed with ultrapure water, dried with nitrogen, annealed at 550°C for 1 hour and coated with an anti-adhesive coating (NXT-110, Nanonex NXT-110). The resulting mold was used to imprint glass coverslips. Glass coverslips (22x22 mm, Menzel Glasser) were cleaned in freshly prepared piranha solution for 15 min (30% $H_2O_2$; 70% $H_2SO_4$), copiously rinsed with water and spin-coated with 40nm of Poly(methyl methacrylate) (PMMA, 494K) and baked for 1 min at 180°C. Then, the PMMA coated coverslips were imprinted using a commercial nanoimprinting tool (NX-B100, Nanonex) with the following parameters: temperature 180℃, pressure 480 psi for 4 minutes. 15nm of Ti was then deposited at an angle (60°) on the imprinted samples to ensure that the mask covered the top PMMA surface but not the bottom of the imprinted features. Then, the exposed PMMA was etched by oxygen plasma for 1 minutes (Corial 200 IL, plasma conditions), the pressure in the chamber was maintained at 10 psi and the oxygen flow was 10 SCCM. The samples were coated with two metals at opposite angles by sequential evaporation of Ti/Au (1 nm/5 nm) and Ti/Cr (5 nm/3 nm). The interdot separation was obtained by changing the angles of evaporation of the metals. Finally, liftoff of the PMMA in boiling acetone was used to obtain array of nanodot heterodimers.

*Preparation of the bimetallic nano-islands with systematically controlled metal ratios*

A silicon wafer (<100> orientation, P-type, 5-10 Ω.cm, thickness 380±25 µm) with a dry oxide layer of 100 nm was cut into 2 by 2 cm samples. The cleaning procedure consisted of sonication in acetone for 10 minutes, cleaning in piranha solution for 10 minutes (30/70 $H_2O_2$:$H_2SO_4$) followed by copious rinsing in DI water and drying with nitrogen. The samples were cleaned in plasma for 2 minutes and then immediately was coated with the resist. The coating procedure consisted of covering the samples with 200 µL of PMMA dissolved in anisole and spin-coating at 4000rpm for 40 seconds followed by baking on a hot plate for 2 minutes at 140 °C. The obtained film thickness was ~40 nm as measured by ellipsometry. An E-beam lithography process was carried using a Raith e-line machine following a CAD model of 45 fields. Each field is made of a matrix of rectangles with increased angular orientation. The writing was made with 10 KeV beam using a dose of 70 $\mu C/cm^2$ and a current of 130pA. The samples were developed in MIBK-isopropanol (1:3 v/v) for 1.5 minutes and then were washed in isopropanol and dried with nitrogen. Then, the wet etching was carried out on the developed samples with a 10% HF buffer solution for 45 seconds in order to create an undercut to the features inside the oxide layer. Using a simulation by PVSyst6.0.1 and linear optics calculations using MATLAB, evaporating parameters were calculated so the feature orientation will encode a predefined Au/$TiO_2$ deposition ratio. The metal deposition was carried using a VST E-Gun metal evaporator. The samples were attached to a tilted sample holder (22°) and placed in the UHV chamber. The evaporation process first consisted of 3 nm of Ti and 6 nm of Au, followed by a 90° rotation of the sample holder and then evaporating 6 nm of Ti and 3 nm of Cr. The anisotropic double evaporation on the tilted features will create a controlled gradient in the ratios between the two metals. The lift-off process was carried using warm (80 °C) NMP for 4 hrs. and then rinsing in DMSO for few more hrs., followed by rinsing in ethanol and water and drying under a stream of nitrogen. After lifting off the resist, the bimetallic nano-islands were imaged by scanning electron microscopy.

**2) Selective chemical biofunctionalization of the samples**

*Chemicals*

Analytical reagent grade sodium hydroxide, acetone, isopropanol, toluene and absolute ethanol were purchased from BioLab Ltd. (Israel) and used as is. Elga PureLab Flex system was used to obtain ultrapure water. Silane terminated poly(ethylene glycol) (PEG-silane; Mw=5000) was purchased from Nanocs (Zotal, Israel). N-Methyl-2-pyrrolidone (NMP), dimethyl sulfoxide (DMSO), 6-amino-1-hexanethiol ($H_2N$-$C_6$-SH), 5(6)-carboxytetramethylrhodamine N-succinimidyl ester (TAMRA-NHS), acetic acid, Cr etchant standard, 11-phosphonoundecanoic acid ($PO_3$–$C_{11}$–COOH), nitrilotriacetic acid-terminal SAM formation reagent (thiol–NTA), *N*-(3-dimethylaminopropyl)-*N′*-ethylcarbodiimide hydrochloride (EDC), *N*-hydroxysuccinimide (NHS), 2-(*N*-morpholino)ethanesulfonic acid

(MES), *O*-(2-aminoethyl)-*O′*-[2-(biotinylamino)ethyl]octaethylene glycol ($NH_2$–$PEG_8$–biotin), and nickel(II) chloride hexahydrate ($NiCL_2 \cdot 6H_20$) were all purchased from Merck (Israel).

*Reagents for Biofunctionalization*

Dulbecco's phosphate-buffered saline (PBS), skim milk, paraformaldehyde, and Tween20 were purchased from Merck. NeutrAvidin Oregon Green 488-conjugated and anti-mouse Alexa Fluor 555 were obtained from Life Technologies (Rhenium, Israel). Biotinylated anti-human KIR2DL1 mAb (biotin α-KIR2DL1) was purchased from Creative Diagnostics (USA). Human MICA protein, His tag (His-MICA), was obtained from SinoBiological (China). Mouse anti-human MICA mAb (α-MICA) was purchased from Abcam (Biotest, Israel). Allophycocyanin labelled anti-human Lysosome-Associated Membrane Protein 1(APC α-CD10a) was obtained from Biolegend (Enco, Israel). Dako Fluorescent Mounting Medium S3023 was purchased from Agilent (USA).

*Chemical Functionalization of the nanodot heterodimers*

To obtain a ternary chemistry on $TiO_2$/Au bimetallic nanoarrays on a $SiO_2$ background, the modification procedure is based on previous work (3, 4), with important modifications. First, to ensure the optimal removal of the resist as well as surface impurities, chromium protected nanopatterned surfaces were alternatively cleaned for one week in DMSO and NMP under mild agitation interspersed with brief sonication. Samples were then copiously rinsed with ethanol and water, dried under a stream of nitrogen and plasma cleaned for 1 min. (Harrick Plasma, USA). Immediately after plasma cleaning, the $SiO_2$ background was passivated by immersing the samples 48 hrs. in a toluene solution containing 1:10000 w/v PEG-silane and 0.1% v/v acetic acid. After this, the surfaces were abundantly rinsed with toluene, acetone, isopropanol, ethanol and water respectively, followed by etching of the protecting Cr layer and rinsing in water. Then, to functionalize $TiO_2$, the surfaces were immersed in a 5 mM aqueous solution of $PO_3$–$C_{11}$–COOH for 24 h. To facilitate the dissolution of $PO_3$–$C_{11}$–COOH in water, a catalytic amount of NaOH was also added. Upon complete reaction time, the samples were directly placed in an oven at 120 °C for 24 h without rinsing. The surfaces were then copiously rinsed with water and immersed for 1 h in an aqueous solution of EDC, NHS, and MES at the concentrations of 0.2, 0.1, and 0.1 M, respectively followed by rinsing with water. $TiO_2$ was then functionalized with biotin by overnight incubation in a 1 mM aqueous solution of $NH_2$–$PEG_8$–biotin. The samples were then copiously rinsed with water and ethanol. Au was functionalized by reacting overnight in a 0.2 mM ethanolic solution of thiol–NTA, followed by generous rinsing with ethanol and water.

*Biofunctionalization of the nanodot heterodimers*

The samples were blocked for 30 min at 37 °C in PBS with 5% skim milk, followed by 30 min incubation in 25 μg/mL NeutrAvidin Oregon Green 488 in PBS with 5% skim milk at room temperature. The surfaces were rinsed 3 × 5 min in PBS with 0.1% Tween20. The samples were then immersed overnight at room temperature in a solution containing 2 μg/mL of biotin α-KIR2DL1 in PBS with 5% skim milk, followed by rinsing 3 × 5 min in PBS with 0.1% Tween20. The surfaces were then incubated for 2 h in nickel(II) chloride, followed by rinsing in water and incubated overnight at 4 °C in His-MICA at a concentration of 2 μg/mL in PBS. Finally, the samples were rinsed 2 × 5 min in PBS with 0.1% Tween20 plus once with neat PBS and stored in PBS until further use. For cell studies, freshly prepared samples were used as is.

*Indirect immunofluorescence labelling of trifunctional nanodot heterodimers*

To confirm the biofunctionalization by fluorescence, the surfaces were incubated overnight in ɑ-MICA at 4 °C in PBS with 5% skim milk (1:100 v/v) followed by rinsing 3 × 5 min in PBS with 0.1% Tween20. Samples were incubated 1 hr. at 37 °C anti-mouse Alexa Fluor 555 in PBS with 5% skim milk (1:40 v/v) followed by rinsing at least overnight in PBS with 0.1% Tween20 in the dark. Finally, surfaces were rinsed once in water, mounted with Dako Fluorescent Mounting Medium and imaged.

**3) Cell studies**

*Primary NK Cell Purification*

pNK cells were purified from peripheral blood of healthy, adult, volunteer donors, recruited by written informed consent, as approved by the Institutional Review Board Ben-Gurion University of the Negev. The cells were isolated using a human negative selection-based NK isolation kit (RosetteSep, Miltenyi Biotec). The purified NK cells were then cultured in a stem cell serum-free growth medium (CellGenix GMP SCGM, 20802-0500) supplemented with 10% heat-inactivated human AB plasma from healthy donors (Sigma, male AB, H-4522), 1% l-glutamine, 1% Pen-Strep, 1% sodium pyruvate, 1% MEM-Eagle, 1% HEPES 1 M, and 300 IU/mL recombinant human IL-2 (PeproTech).

*NK Cell Activation Studies*

Cultured pNK cells were seeded onto the surfaces in growth medium containing <2% serum, 50 units of Il-2 which was supplemented with APC ɑ-CD10a (1:1000 v/v) and left to adhere for 3–4 h. The surfaces were then rinsed twice in PBS to remove the nonadherent cells, followed by fixing the adherent cells with 4% paraformaldehyde (PFA) and then direct staining with Alexa Fluor 555 phalloidin without permeabilization to prevent damage to the cell membrane. Finally, the nuclei were stained by mounting the samples with ProLong Gold antifade reagent containing DAPI (both from Life Technologies).

*Microscopy*

The characterization of the surfaces as well as NK cell adhesion and activation were performed using a Nikon Ti2e epifluorescence microscope and quantified using the Fiji imaging software (https://fiji.sc). For ɑ-CD107a quantification of fluorescence intensity, exposure time, and magnification were not changed between samples. We quantified the degree of activation, which is proportional to the intensity of the APC ɑ-CD107a signal (5).

*Statistics*

At least 10 fields at 20× magnification on each surface were analyzed. The data were averaged for each experiment, and the experiments were performed three times. Statistical analysis was performed by analysis of variance, and Tukey's multiple comparison *post hoc* test was also performed using the Prism software (GraphPad Software Inc., USA). The results were considered to be significantly different for $p < 0.05$.

**4) Determination of the separation between the nanodot heterodimers**

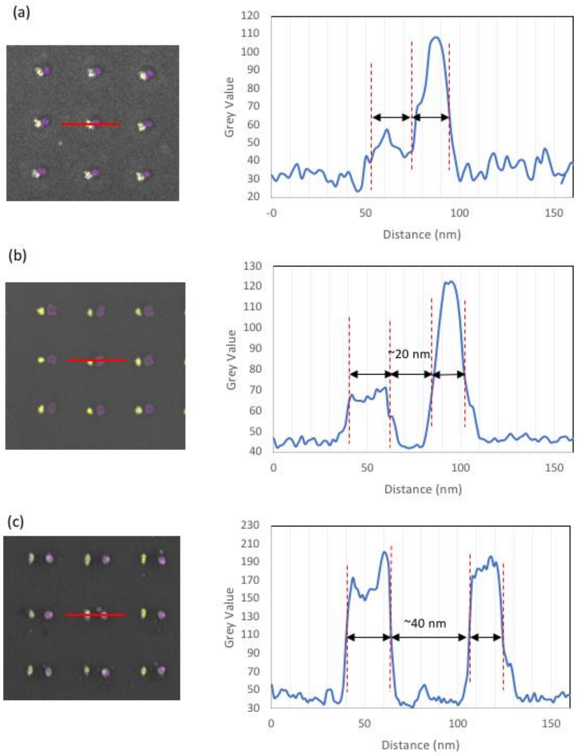


Fig. S1. Dimensional analysis of dimers consisting of Au and Ti nanodots with the gap of (a) 0 nm, (b) 20 nm, and (c) 40 nm. The dimers were analyzed by cross-section profiling using ImageJ software. The feature dimensions were assessed based in the Full Width Half Maximum (FWHM) of the peak profiles, and were found to be in the range of ~15-20 nm.

**5) Quantification of biofunctionalization by surface plasmon resonance (SPR)**

Surface plasmon resonance confirmed the effective grafting of a biotinylated anti-KIR2DL1 to a streptavidin modified sensorchip with a surface density of 1 molecule per 210 $nm^2$ assuming that the immobilization is homogeneous onto the surface. In addition, the attachment of a histidine tagged MICA to a nickel(ii)-nitrilotriacetic functionalized sensorchip at a surface density of 1 molecule per 190-230 $nm^2$ was demonstrated.

Bare gold SPR sensorchips from Xantec Bioanalytics (Germany) were used for SPR experiments. sensor chips were functionalized with either biotin or NTA accordingly. Analyses were performed on a Biacore T200 instrument. After introduction of the mounted chips, samples were flushed with PBS. Homogeneity of the surface modification was assessed by comparing plasmon resonance across the 4 available flow cells. Then, one flow cell was chosen and either His-MICA, for $Ni^{2+}$-NTA surfaces, or streptavidin and anti-KIR2DL1, for biotin surfaces were immobilized. Ligands or antibodies were injected at various concentrations. A low flow rate was set at 5-10 µl/min for all samples.

*His-MICA on Ni-NTA modified gold Sensorchips*

The real-time evolution of the SPR response in resonance units (RU) as a function of His-MICA injection is shown in figure S2.

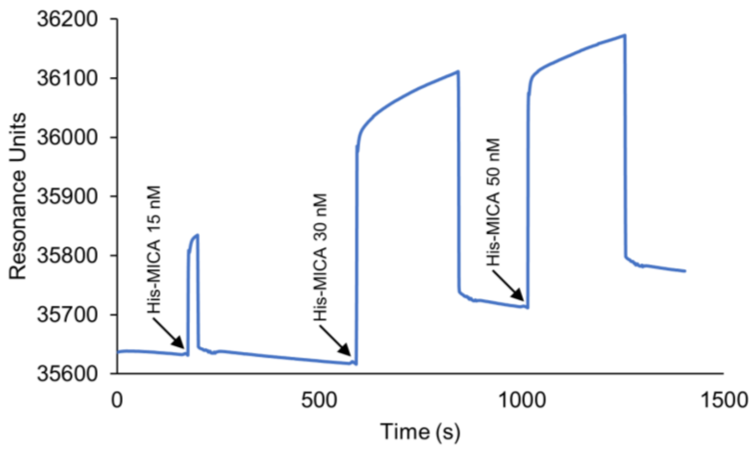


Figure S2. Changes in SPR response as His-MICA is injected onto a Ni-NTA modified sensor chip at concentration ranging from 15 to 50 nM.

Coupling of His-MICA was visible by the stepwise increase in RU following injection. The calculated rate of immobilization was 50 RU/min. However, the kinetics appeared slow and all the His-MICA would have been consumed before maximum loading capacity would have been reached. Instead, it was decided that the sensor chip would be incubated overnight in His-MICA. The method was, for example, used to functionalized surfaces for PM-IRRAS measurements. The sensor chip would then be introduced in the machine and the initial resonance would be recorded (figure S3). Changes in RU at $t_0$ in Fig. S2 and $t_0$ in Fig. S3 (ΔRU) would correspond to the amount of surface grafted His-MICA, where ΔRU = 1000 : 1 ng/mm² as per Biacore.

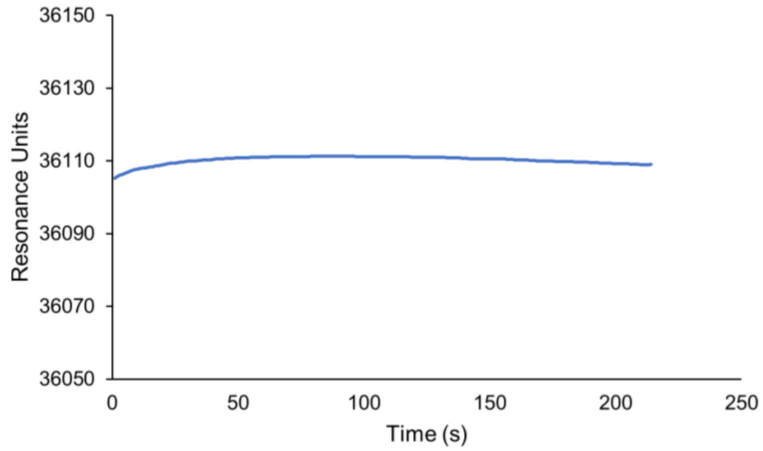


Figure S3: SPR output after overnight incubation of the $Ni^{2+}$-NTA sensor chip in 2 µg/mL His-MICA.

A value of ΔRU = 36105 – 35638 = 467 was found. This equates to 467 pg/mm². As per the supplier, the molecular weight of His-MICA is 55-65 kDa. Therefore, the maximum loading capacity was found to be 7.2-8.5 fmol/mm². This value is substantially lower than the 130 fmol/mm² which were found using immunofluorescence (4). This may be due to a higher amount of non-specific interactions occurring during the immunofluorescent staining of the surfaces. Finally, the loading capacity can be translated in surface density and it was found that there is one His-MICA molecule per 190-230 nm².

*Streptavidin and biotinylated anti-KIR2DL1 on biotin modified gold Sensorchips*

The real-time evolution of the SPR response in resonance units (RU) as a function of the sequential injection of streptavidin followed by a biotinylated anti-KIR2DL1 is shown in figure S4.

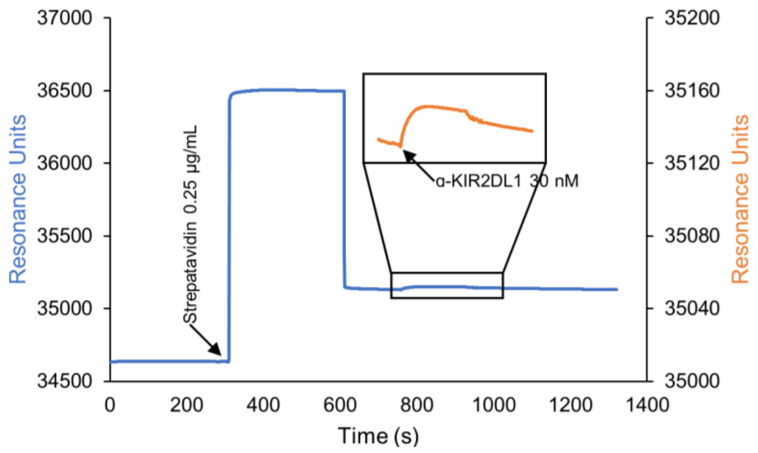


Figure S4. Changes in SPR response as streptavidin (0.25 µg/mL), followed by biotinylated anti-KIR2DL1 (30 nM) are sequentially injected onto a biotin functionalized sensor chip. Inset shows the injection of anti-KIR2DL1.

Coupling of streptavidin to the biotin modified surface was confirmed by the 507 RU increase in the SPR response between 300 and 620 s (Fig. S4). Considering that for streptavidin MW = 52.8 kDa, the loading capacity of our surfaces was found to be 9.6 fmol/mm². Then, from 760 to 925 s, the further increase in resonance, resulting from the injection of biotinylated anti-KIR2DL1 confirms its successful grafting to the streptavidin functionalized sensor chip. In this case, For the attachment of biotinylated anti-KIR2DL1, the value ΔRU = 26.5 corresponds to a loading capacity of 7.8 fmol/mm². Assuming that immobilizations are homogeneous on the surface, the loading capacity found here implies that (i) considering a MW=150 kDa, there is one biotinylated anti-KIR2DL1 per 210 nm² and that (ii) there are 0.81 biotinylated anti-KIR2DL1 for every streptavidin on our surfaces. Although, streptavidin has 4 biotin binding sites, at least one site will be devoted to coupling onto the biotin functionalized surfaces. The current result indicates that roughly only one site is available for further grafting of biotinylated molecules.

### 6) Chemical modification of $TiO_2$ and Au is confirmed by Polarization Modulation Infrared Reflection Absorption Spectroscopy (PM-IRRAS)

For gold surfaces, 15 nm of titanium and 200 nm of gold were successively evaporated onto 24x24 mm coverslips. For titanium surfaces, 15 nm of titanium, 200 nm of gold and again 15 nm of titanium were evaporated onto 24x24 mm coverslips. Samples were left in air to allow the formation of a native $TiO_2$ passivating layer. The $TiO_2$ and gold surfaces were functionalized with biotin and $Ni^{2+}$-NTA respectively. Importantly, to demonstrate the orthogonality of our chemical modification, both surface types underwent the entire chemical modification to obtain the previously described trifunctional nanodot heterodimers .

PM-IRRAS spectra were recorded on a ThermoNicolet Nexus 670 FTIR spectrometer at a resolution of 4 $cm^{-1}$, by coadding several blocks of 1500 scans (30 minutes acquisition time). All spectra were collected in a dry-air atmosphere after 30 min of incubation in the chamber. Experiments were performed at an incidence angle of 75° using an external homemade goniometer reflection attachment (6). The infrared parallel beam was directed out of the spectrometer with an optional flipper mirror and made slightly convergent with a first $BaF_2$ lens. The IR beam passed through a BaF2 wire grid polarizer (Specac) to select the p-polarized radiation and a ZnSe photoelastic modulator (PEM, Hinds Instruments, type III) which modulates the polarization of the beam at a high fixed frequency (74 KHz) between the parallel (p) and perpendicular (s) linear states. After reflection on the sample, the double modulated (in intensity and in polarization) infrared beam was focused with a second ZnSe lens onto a photovoltaic MCT detector (Kolmar Technologies, Model KV104) cooled at 77 K. In all experiments, the PEM was adjusted for a maximum efficiency at 2500 $cm^{-1}$ to cover the mid-IR range in only one spectrum. For calibration measurements, a second linear polarizer (oriented parallel or perpendicular to the first preceding the PEM) was inserted between the sample and the second ZnSe lens. This procedure was used to calibrate and convert the PM-IRRAS signal in terms of the IRRAS signal (i.e., 1–Rp(d)/Rp(0) where Rp(d) and Rp(0) stand for the p-polarized reflectance of the film/substrate and bare substrate systems, respectively) (7).

*Biotin-PEG-SH monolayer on Au surfaces*

PM-IRRAS spectra are usually acquired on gold substrates. To demonstrates the feasibility of PM-IRRAS on titanium we first characterized biotin terminated poly(ethylene glycol) SAM on gold. The PM-IRRAS spectrum of the biotin functionalized gold surfaces is shown in Fig. S5.

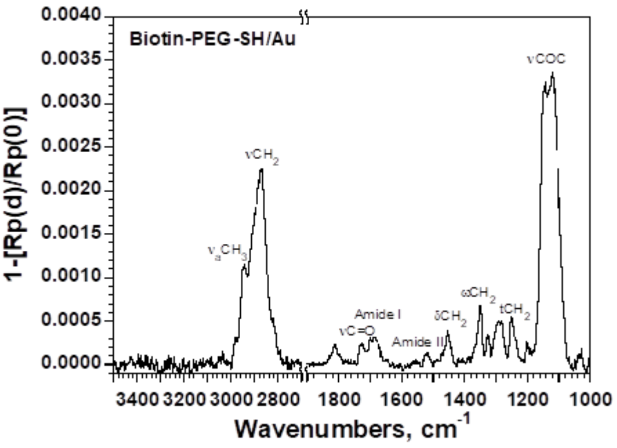


Fig. S5. PM-IRRAS spectrum of biotin terminated SAMs on gold with assignment of the characteristic bands.

The presence of νC=O band assigned to biotin, amide I and amide II bands from the amide groups and $\nu CH_2$, $\delta CH_2$, $\omega CH_2$, $tCH_2$ and νCOC from the poly(ethylene glycol) backbone confirms that the gold surface is functionalized by biotin ligands. The presence of $\nu_a CH_3$ band stemming from the distal methoxy PEG used as a spacer with the monolayer to limit steric hindrance of the biotin moieties confirms that mixed SAM are used here.

*Biotin-$PO_3$ monolayer on $TiO_2$ surfaces*

The PM-IRRAS spectrum of a Biotin terminated alkylphosphonic acid SAM on $TiO_2$ surface is shown in Fig. S6.

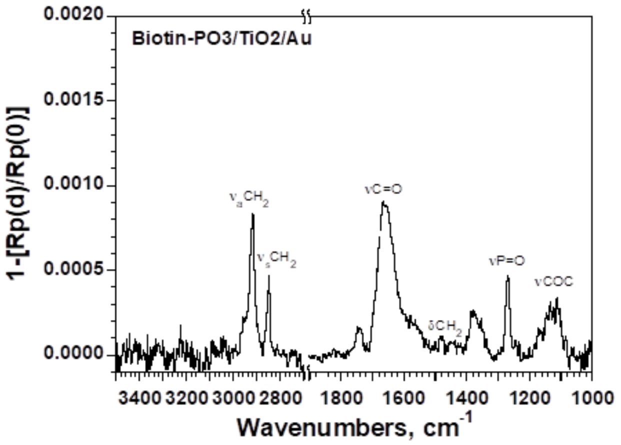


Fig. S6. PM-IRRAS spectrum of biotin functionalized $TiO_2$ with assignment of characteristic bands.

The νP=O band confirms that the alkyl phosphonic base layer used to graft biotin to the $TiO_2$ surface is present. However, the intensity of PEG bands is very low. In contrast, the intensity of the νC=O band relative to the biotin moiety is higher than for the gold surface.

*Ni-NTA-SH monolayers on gold surfaces*

The PM-IRRAS spectrum of nickel(ii) NTA functionalized Au surface is shown in Fig. S7.

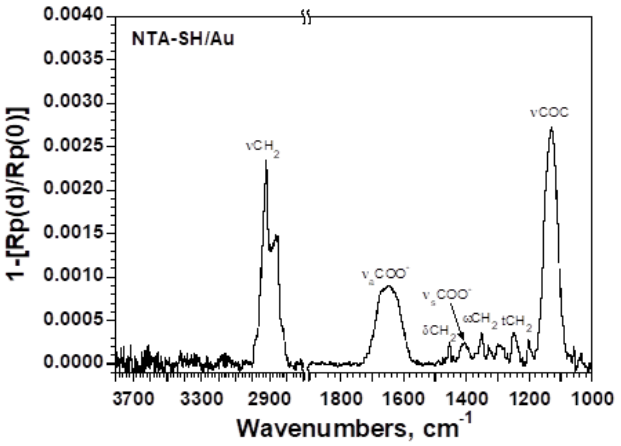


Fig. S7. PM-IRRAS spectrum of a Ni-NTA terminated SAM on gold, with assignment of characteristic bands.

The $\nu_aCOO^-$ and $\nu_sCOO^-$ bands reveal the presence of carboxylate moieties that are characteristic of a NTA complex with a chelated metal ion, $Ni^{2+}$-NTA complex in our case. The $\nu_aCH_2$ and $\nu_sCH_2$ bands correspond to the alkyl chains and $\nu CH_2$, $\delta CH_2$, $\omega CH_2$, $tCH_2$ and νCOC bands correspond to the poly(ethylene glycol) backbone which both constitute the Au-bound base layer. The gold surface is functionalized by $Ni^{2+}$-NTA complex.

**7) Confirmation that the ratio between the two metals is controlled by the feature orientation according to the PVSyst6.0.1 simulation.**

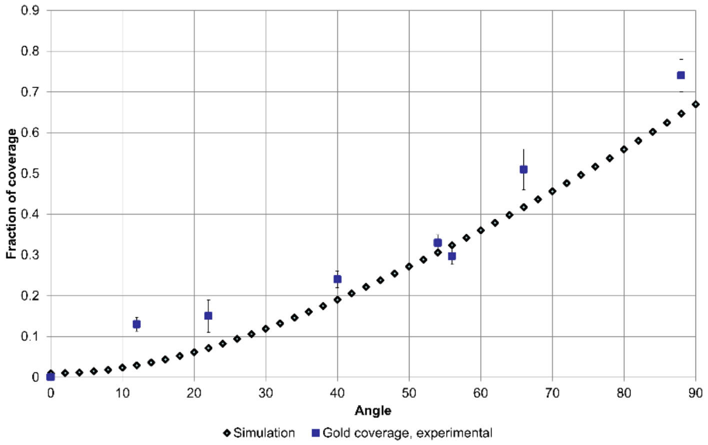


Fig. S8. Gold coverage as a function of feature orientation. Experimental data obtained by SEM (blue squares) is compared to the simulation by PVSyst6.0.1 (black rhombi).

### 8) Estimation of the ligand length

The length of NTA linker to Au was assessed using Chemsketch software, and was found to be 4.22 nm long. The crystalline structure of the MICA-NKG2D complex is 10.18 nm long(8). Also, the crystalline structure of NKG2D alone is 3.61 nm long(9). So, the approximate length of MICA construct, without NKG2D, is 10 nm.

The length of biotin linker to Titanium was assessed using Chemsketch software, and was found to be 6.54 nm. The crystalline structure is 4,35 nm long(10). The crystal structure of immunoglobulin is 10.61 nm long(11). So, the approximate length of KIR2DL1 construct is 22 nm.

### 9) Calculation of the bond affinity

In order to understand how the spatial coordination of ligands for NKG2D and KIR2DLI mediates the CD107a expression, we first calculate the affinity of bonds between the two proteins and their respective ligands, if appearing as an isolated pair (Fig. S9).

Due to cooperative effects these bonds are more stable together than each individually, since they share the stresses produced by the cellular membrane. However, due to a complex profile of membrane mediated forces on bonds in the membrane (12, 13), these effects are strongly dependent on the separation between the bonds. Using the previously dependent formalism (12, 13), it is easy to show that the binding and unbinding rates $K_{\mathrm{on}}$ and $K_{\mathrm{off}}$ adopt the form:

$$K_{\mathrm{on}} = k_0 \frac{\sqrt{\lambda\alpha^2}}{\sqrt{2\pi(1+\lambda\sigma)}} \exp\left[-\frac{\lambda\{\bar{h}-(\alpha+l_0)\}^2}{2(1+\lambda\sigma)}\right] \quad \text{(S1)}$$

$$K_{\mathrm{off}} = k_0 \exp\left[\frac{\lambda\alpha}{2}\left[2\left(\bar{h}-l_0\right)+\alpha(\lambda\sigma-1)\right]-\epsilon_b\right]. \quad \text{(S2)}$$

Here, $k_0$ is the intrinsic binding rate, $l$ is the stiffness of the receptor-ligand bond, $\bar{h}$ and $\sigma$ are the average position and fluctuation amplitude of the cell membrane at the position of the bond, $\alpha$ is the size of the binding pocket and $\epsilon_b$ is the intrinsic binding affinity. The binding affinity of bond can be determined by

$$E_b = \log\frac{K_{\mathrm{on}}}{K_{\mathrm{off}}}. \quad \text{(S3)}$$

In order to calculate the binding affinity for the current problem of two bonds, we first consider a formation process at **r** of a short bond with rest length $l_S$ and stiffness $\lambda_S$ in a presence of an existing long bond located at **0** with rest length $l_L$ and stiffness $\lambda_L$. Therefore, the membrane mean height and fluctuation amplitude can be given by (12, 13)

$$\bar{h}(\mathbf{r}) = -\frac{\pi(l_L-h_0)\mathrm{kei}(q_0|\mathbf{r}|)}{\frac{\pi^2 g}{4\xi_\perp^2}+4\mathrm{kei}^2(q_0|\mathbf{r}|)} \qquad \text{(S4)}$$

$$\sigma(\mathbf{r}) = \frac{g}{\frac{g}{\xi_\perp^2}+\frac{16}{\pi^2}\mathrm{kei}^2(q_0|\mathbf{r}|)}, \qquad \text{(S5)}$$

with $g = \frac{1}{\lambda_L} - \frac{4\xi_\perp^2\mathrm{kei}(0)}{\pi} - \frac{16\xi_\perp^2\mathrm{kei}^2(q_0|\mathbf{r}|)}{\pi^2}$. Here, $\xi_\perp^2$ and $h_0$ stand for the fluctuation amplitude and mean height of membrane in unbound state, kei(·) and $q_0$ denote the Kelvin-function and characteristic model, respectively. Then, inserting Eqs. (S4, S5) into Eq. (S1) yields the binding rate of short bond as

$$K_{\mathrm{on}}^S = k_0\frac{\sqrt{\lambda_S\alpha^2}}{\sqrt{2\pi\left(1+\frac{\lambda_S g}{\frac{g}{\xi_\perp^2}+\frac{16}{\pi^2}\mathrm{kei}^2(q_0|\mathbf{r}|)}\right)}}\exp\left[-\frac{\lambda_S\left\{\frac{\pi(l_L-h_0)\mathrm{kei}(q_0|\mathbf{r}|)}{\frac{\pi^2 g}{4\xi_\perp^2}+4\mathrm{kei}^2(q_0|\mathbf{r}|)}+(\alpha+l_S)\right\}^2}{2\left(1+\frac{\lambda_S g}{\frac{g}{\xi_\perp^2}+\frac{16}{\pi^2}\mathrm{kei}^2(q_0|\mathbf{r}|)}\right)}\right]. \qquad \text{(S6)}$$

Once the short bond is closed in a presence of a long bond has been obtained, we further calculate the dissociation rate of the short bond. In the case of formation of both two bonds, the related membrane mean height and fluctuation amplitude at the position of short bonds can be given by (12, 13)

$$\bar{h}(\mathbf{r}) = -\frac{\frac{4}{\pi}\left[\left(G_{11}^{-1}+G_{12}^{-1}\right)(l_S-h_0)\mathrm{kei}(0)+\left(G_{21}^{-1}+G_{22}^{-1}\right)(l_L-h_0)\mathrm{kei}(q_0|\mathbf{r}|)\right]}{8\sqrt{\kappa\gamma}+\frac{16}{\pi^2}\left[G_{11}^{-1}\mathrm{kei}^2(0)+\left(G_{12}^{-1}+G_{21}^{-1}\right)\mathrm{kei}(0)\mathrm{kei}(q_0|\mathbf{r}|)+G_{22}^{-1}\mathrm{kei}^2(q_0|\mathbf{r}|)\right]}, \qquad \text{(S7)}$$

$$\sigma(\mathbf{r}) = \left\{\frac{1}{\xi_\perp^2} + \frac{16}{\pi^2}\left[G_{11}^{-1}\mathrm{kei}^2(0) + \left(G_{12}^{-1} + G_{21}^{-1}\right)\mathrm{kei}(0)\mathrm{kei}(q_0|\mathbf{r}|) + G_{22}^{-1}\mathrm{kei}^2(q_0|\mathbf{r}|)\right]\right\}^{-1}, \qquad \text{(S8)}$$

with the coupling matrix $G_{ij}$ defined as

$$G_{11} = \frac{1}{\lambda_S} - \frac{4\xi_\perp^2\mathrm{kei}(0)}{\pi} - \frac{16\xi_\perp^2\mathrm{kei}^2(0)}{\pi^2},\ G_{12} = G_{21} = -\frac{4\xi_\perp^2\mathrm{kei}(q_0|\mathbf{r}|)}{\pi} - \frac{16\xi_\perp^2\mathrm{kei}(0)\mathrm{kei}(q_0|\mathbf{r}|)}{\pi^2},$$
$$G_{22} = \frac{1}{\lambda_L} - \frac{4\xi_\perp^2\mathrm{kei}(0)}{\pi} - \frac{16\xi_\perp^2\mathrm{kei}^2(q_0|\mathbf{r}|)}{\pi^2}.$$

Subsequently, inserting Eqs. (S7, S8) into Eq. (S2) yields the relevant dissociation rate of the short bond as

$$K_{\mathrm{off}}^S = k_0\exp\left\{\frac{\lambda_S\alpha}{2}\left[2\left(-\frac{\frac{4}{\pi}\left[\left(G_{11}^{-1}+G_{12}^{-1}\right)(l_S-h_0)\mathrm{kei}(0)+\left(G_{21}^{-1}+G_{22}^{-1}\right)(l_L-h_0)\mathrm{kei}(q_0|\mathbf{r}|)\right]}{\frac{1}{\xi_\perp^2}+\frac{16}{\pi^2}\left[G_{11}^{-1}\mathrm{kei}^2(0)+\left(G_{12}^{-1}+G_{21}^{-1}\right)\mathrm{kei}(0)\mathrm{kei}(q_0|\mathbf{r}|)+G_{22}^{-1}\mathrm{kei}^2(q_0|\mathbf{r}|)\right]} - l_S\right) + \alpha\left(\frac{\lambda_S}{\frac{1}{\xi_\perp^2}+\frac{16}{\pi^2}\left[G_{11}^{-1}\mathrm{kei}^2(0)+\left(G_{12}^{-1}+G_{21}^{-1}\right)\mathrm{kei}(0)\mathrm{kei}(q_0|\mathbf{r}|)+G_{22}^{-1}\mathrm{kei}^2(q_0|\mathbf{r}|)\right]} - 1\right)\right] - \epsilon_b\right\}. \qquad \text{(S9)}$$

Submitting Eqs. (S6, S9) into Eq. (3) gives the binding affinity of the short bond in presence of a long bond. Similarly, we also can calculate the binding affinity of a long bond in presence of a short bond as a function of membrane correlation length as shown in Fig. S9, which only requires the swap of indices L and S in Eqs S1-S9.

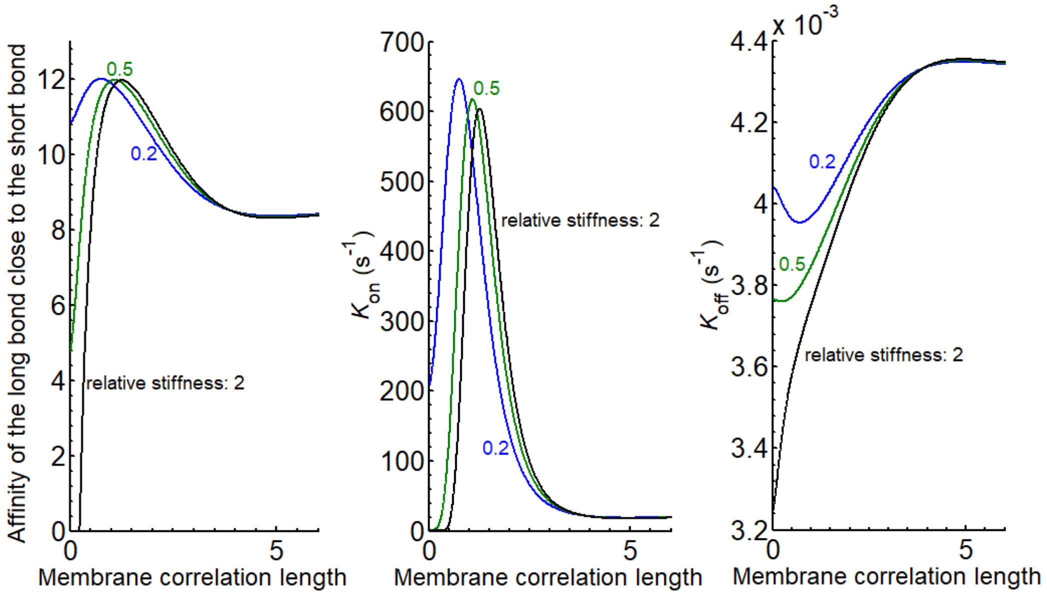


Fig. S9. Affinity and reaction rates of the long bond close to the short bond as a function of membrane correlation length for $h_0 = 70$ nm, $\xi_\perp^2 = 51$ nm$^2$, $q_0 = 0.0064$ nm$^{-1}$, $l_L = 50$ nm, $l_S = 30$ nm, $\alpha = 1$ nm, $k_0 = 10000$ s$^{-1}$ $\epsilon_L = \epsilon_S = 15\ k_BT$ and $\lambda_L = 2 \times 10^5\ k_BT/\mu\text{m}^2$.

**10) Monte Carlo simulations**

Based on a recently published protocol (12–14), we also propose the relative Monte Carlo simulation on membrane adhesion via two types of receptor-ligand bonds. In detail, the membrane and substrate are discretized into square lattices with lattice constant $a$ and decorated with receptors and ligands, respectively. Each receptor or ligand occupies a single site. In simulation system, the ligands are regularly fixed on the surface of substrate. Particularly, the gap between one type of ligand and another type of ligand is given value. Meanwhile, the neighbours of same type of ligands keep a constant gap as 200 nm. Initially, the receptors are randomly positioned on the membrane surface. The periodic boundary conditions are considered here. A contact zone with a fixed radius $R$ is located in the middle of the simulation box, while receptors on the free region of membrane keep constant density, which means it can provide the reservoir reconstructing the appropriate statistical ensemble. The mobility of freely diffusive ligands is treated as a random walker with time step $\Delta t = a^2/D$. Here, $D$ represents the diffusion coefficient of both types of ligands. During each time step $\Delta t$, the stochastic processes of bonds' formation and dissociation would happen. The simulations are performed until equilibrium states are reached. The representative values of the related parameters in the Monte Carlo simulations are shown in Table S1

Table S1 Values of the system parameters

| Parameter | Symbol | Value |
|---|---|---|
| Ligand separation – cell size | $a$ | 5, 20 and 40 nm |
| Number of cells | $N$ | 500×500 |
| Radius of contact zone | $R$ | 200 number of cells |
| Diffusion coefficient of receptors | $D$ | 0.002 $\mu m^2/s$ |
| Normalized density of receptors | $\rho_L^r = \rho_S^r$ | 0.2 |
| Mean height | $h_0$ | 70 nm |
| Fluctuation amplitude | $\xi_\perp^2$ | 51 $nm^2$ |
| Characteristic model of membrane | $q_0$ | 0.0064 $nm^{-1}$ |
| Interaction range | $\alpha$ | 1 nm |
| Intrinsic reaction rate | $k_0$ | 10000 $s^{-1}$ |
| Rest length of bonds | $l_L$ | 50 nm |
| | $l_S$ | 30 nm |
| Stiffness of bonds | $\lambda_L$ | $2\times10^5\ k_BT/\mu m^2$ |
| | $\lambda_S$ | Variable |
| Binding energy | $\epsilon_L = \epsilon_S$ | 15 $k_BT$ |

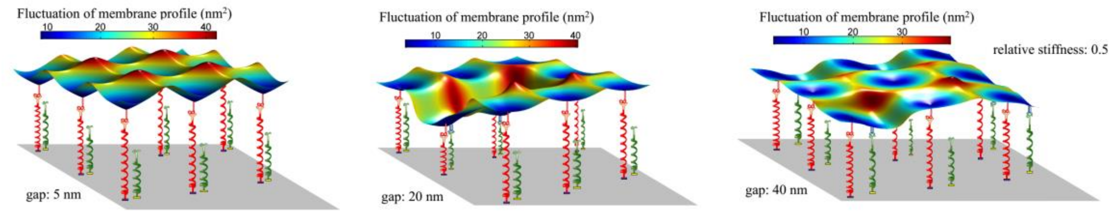


Fig. S10 Snapshot of membrane profile for different gap between red and green binding sites.
